# A Large-Scale Computer-Vision Mapping of the Geometric Structures of Stroboscopically-Induced Visual Hallucinations

**DOI:** 10.64898/2026.02.18.705710

**Authors:** Ethan J. Grove, Trevor Hewitt, Anil K. Seth, Fiona Macpherson, David J. Schwartzman

**Affiliations:** Sussex Centre for Consciousness Science, Department of Engineering and Informatics, University of Sussex, Brighton, England; Program for Brain, and Consciousness, Canadian Institute for Advanced Research (CIFAR), Toronto, Canada; Centre for the Study of Perceptual Experience, University of Glasgow, UK

## Abstract

Visual hallucinations (VHs) occur across psychedelic states and diverse psychiatric and neurological conditions, yet their phenomenology remains difficult to characterise. Empirical research on VHs is hindered by the lack of large-scale phenomenological datasets, which limits both mechanistic accounts and the systematic characterisation of when and how they arise. Stroboscopic light stimulation (SLS) viewed with closed eyes provides a reliable, non-pharmacological method of inducing VHs in healthy populations. These hallucinations typically consist of vivid colours and dynamic geometric patterns that resemble simple VHs described in both psychedelic and clinical contexts, suggesting partially overlapping neural mechanisms. We developed and applied an unsupervised computer-vision pipeline to analyse a large dataset of 10,598 drawings made following exposure to hallucination-inducing SLS. These drawings were produced by attendees of Dreamachine, a large-scale public installation designed to elicit stroboscopically induced visual hallucinations (SIVHs). We extracted feature embeddings with a self-supervised deep vision transformer, then applied dimensionality reduction and density-based clustering to identify recurrent visual motifs in a data-driven manner. The majority of drawings contained geometric forms, consistent with prior observations of simple VHs under SLS. However, we also identified novel and underreported geometric formations, such as concentric squares, crosses, hyperbolic patterns, and other geometries. Our results show how an unsupervised computer-vision pipeline can organise large, openly shared phenomenological datasets into interpretable classes. By mapping the diversity of simple geometric VHs at scale, this work places new constraints on existing theoretical accounts and motivates targeted experimental work linking SLS parameters, neural dynamics, and geometric visual hallucinations.

## 1 Introduction

Visual hallucinations (VHs) are internally generated perceptual experiences that occur in the absence of corresponding external stimuli. They can occur in a wide range of contexts, including psychopathology (e.g., psychosis and schizophrenia) (Oorschot et al. 2012; Sikich 2013; Waters et al. 2014), neurological disorders (Golden and Josephs 2015; O’Brien et al. 2020; Panayiotopoulos 1999), and are often associated with conditions such as Age-Related Macular Degeneration (AMD) and visual field loss (Schadlu et al. 2009; O’Brien et al. 2020; Teunisse et al. 1996). They can also be induced through pharmacological agents (e.g., psychedelic drugs such as mescaline, LSD, or psilocybin) (Makin et al. 2023; Aqil and Roseman 2023), and through non-pharmacological approaches, including stroboscopic light stimulation (SLS). The study of VHs has contributed to understanding the neural and phenomenological organisation of vision and conscious experience, particularly in altered and pathological states (ffytche et al. 1998; Safron et al. 2025; Rieser et al. 2024; Collerton et al. 2016).

Simple VHs are thought to be closely linked to low-level neural processes supporting visual perception (Suzuki et al. 2024; Ermentrout and Cowan 1979; Bressloff et al. 2002). Unlike complex VHs, which involve semantically meaningful content such as faces, people, animals or scenes (Shenyan et al. 2024), simple VHs typically consist of colours, patterns, and dynamic geometric formations (Suzuki et al. 2024; ffytche 2008). The visual qualities of simple VHs appear to be remarkably consistent across contexts and aetiologies, from psychedelics to psychiatric and neurological disorders (Waters et al. 2014; Bressloff et al. 2001; Klüver 1928; Merabet et al. 2004; van Ommen et al. 2019; Panayiotopoulos 1999), making them a promising target for comparative and mechanistic study.

Two frameworks aim to describe the geometry of simple VHs: Klüver’s form constants (Klüver 1928) and neural-field models (Ermentrout and Cowan 1979; Rule et al. 2011; Bressloff et al. 2001). The Klüver framework, which has been particularly influential, is a classification system developed primarily from mescaline reports in the 1920s. This scheme proposes that simple VHs consist of four universal “form constant” geometric patterns: lattices, cobwebs, tunnels, and spirals (Fig. 1), along with combinations and variations of these motifs.

**Figure 1.**
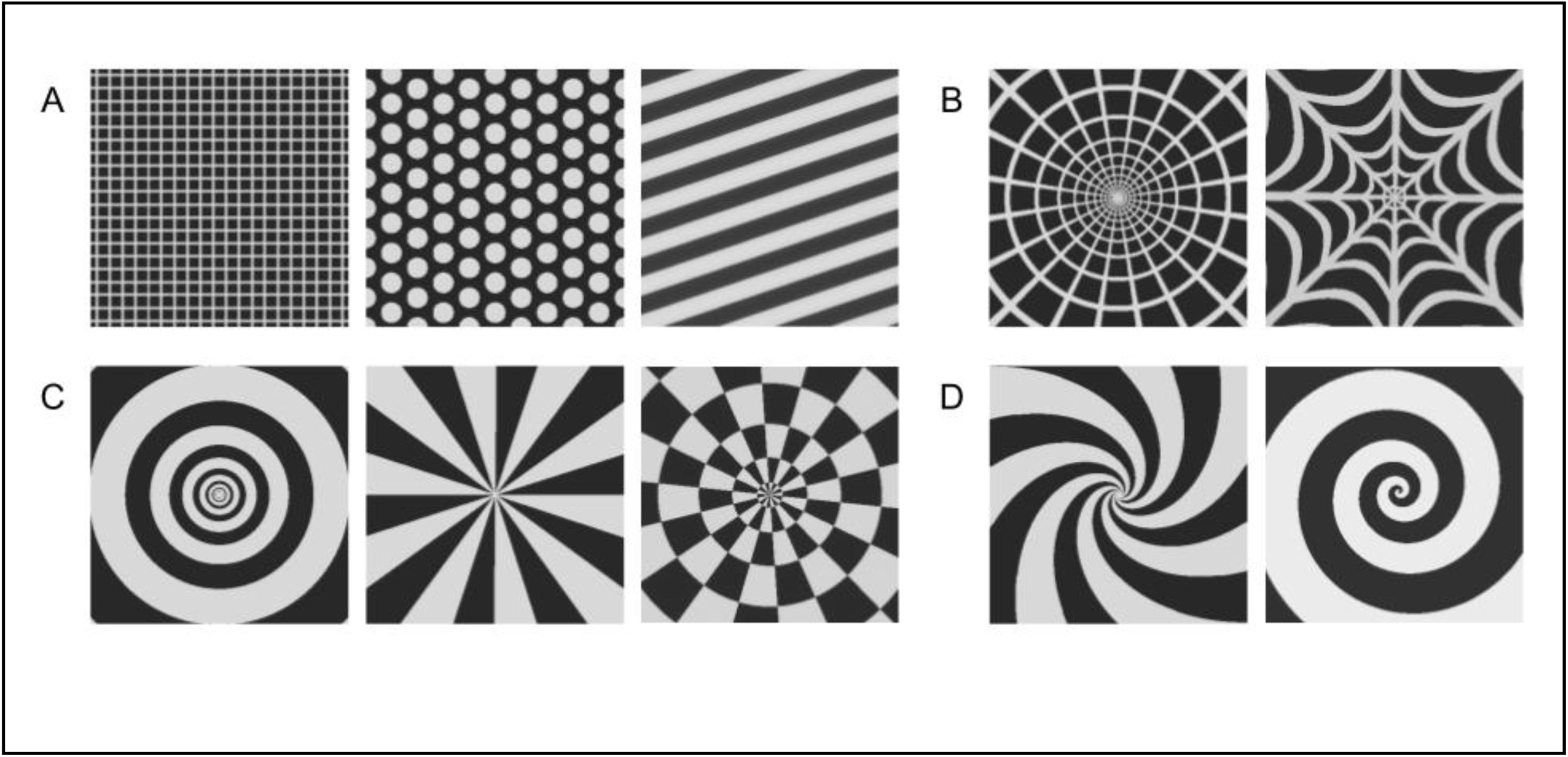
Illustrative examples of Klüver’s four geometric form constants. (A) The lattice, also known as a grating, honeycomb, or checkerboard, illustrates a predominantly translationally symmetric arrangement, characterised by repeating elements spaced along linear axes. (B) The cobweb, (C) the tunnel (or funnel), and (D) the spiral illustrate rotationally symmetric arrangements, characterised by approximately repeating patterns that converge toward a centre, including radial, concentric, and spiral geometries. Translationally symmetric geometries may also exhibit rotational symmetry, whereas predominantly rotationally symmetric geometries typically lack global ranslational structure.

Subsequent mathematical neural-field models sought to account for these form constants by modelling their emergence through the pattern-forming dynamics of primary visual cortex (V1). In these models, cortical patterns with translational symmetry give rise to percepts with rotational symmetry when mapped into visual space via the approximately log–polar retinotopic transform (Fig. S1). This accounts for the cobweb, tunnel, and spiral form constants, which are rotationally symmetric, but does not account for lattices, which are characterised by translational symmetry (see Ermentrout & Cowan, 1979). Here, the term “symmetry” is used in a descriptive phenomenological sense: these patterns exhibit coarse, repeating structure rather than exact mathematical symmetry.

Both Klüver form constants and neural-field models aim to describe the space of possible simple VH geometries. However, this posited space of possible geometries is at best weakly supported by existing evidence (Bressloff et al. 2002), as descriptions and drawings of simple VHs frequently contain geometric structures that fall outside these schemes (Smythies 1959; Mauro et al. 2015; Allefeld et al. 2011). At the same time, these studies typically rely on small samples, heterogeneous report formats, and limited quantitative characterisation of geometry, making it difficult to determine whether the observed range of VH experience can in fact be captured by existing models. Such a determination requires a comprehensive investigation of the phenomenology of VHs. However, research into VHs has generally focused on their correlates, consequences, or diagnostic roles (ffytche 2008; Amaya et al. 2023; Heller et al. 2023; Schwartzman et al. 2019; Bhome et al. 2023; Sikich 2013; Tang and Tang 2020).

Given this context, there is a unique opportunity provided by strobe-light stimulation (SLS). SLS can reliably induce VHs - which can be termed stroboscopically induced visual hallucinations (SIVHs; (Hewitt et al. 2025)). Because SIVH can be readily elicited in non-clinical participants, the use of SLS offers a practical route to large-scale phenomenological data collection.

Adopting this approach, we analysed 10,598 attendee drawings produced following SLS exposure during the Dreamachine public art-science installation, which used SLS to evoke hallucinatory experiences in public audiences (https://dreamachine.world). Leveraging the unprecedented scale of the Dreamachine dataset, we applied an unsupervised computer-vision pipeline, embedding each drawing using a pretrained DINOv2 vision transformer and clustering the resulting features (PCA, UMAP, and HDBSCAN) to identify recurring geometric motifs. This approach yields a data-driven characterisation of the geometric content associated with SIVHs that is independent of existing classification systems or neural-field predictions, enabling direct empirical comparison with these frameworks.

The following sections describe our analytical pipeline, present the resulting clusters, and outline their thematic organisation. We assess how these findings align with, and place constraints on, existing theoretical frameworks based on Klüver form constants and neural-field models. Our results corroborate several core aspects of these frameworks, while also revealing a substantially broader and more diverse range of simple VH geometries than is readily accounted for by current theories, thereby placing new empirical constraints (and opportunities) on future mechanistic models of visual hallucinations.

## 2. Methods

A mixed-methods observational analysis was conducted on 10,598 drawings produced by participants in the Dreamachine installation. We first describe the collection and preprocessing of the drawings, then outline the feature-extraction and clustering procedures used to identify groups of visually similar images, followed by qualitative comparison to existing theoretical and empirical accounts of the content of VHs in the literature. Our mixed-methods approach combines unsupervised computer-vision methods with qualitative interpretation to organise the large-scale dataset into a series of interpretable clusters. All analyses were performed on the dataset publicly released by the event organisers and did not involve any additional data collection.

### Ethics statement

The analyses reported here used a fully anonymised dataset of scanned drawings publicly released by Collective Act Ltd. (CAL), organisers of Dreamachine. The dataset was accessed for research purposes on the 06^th^ July 2022. No demographic or identifying information was available to the authors. According to CREC policies at the University of Sussex, secondary analyses of fully anonymised datasets do not require formal ethical approval. The Dreamachine installation included attendee information and consent procedures overseen by CAL, which informed attendees that their drawings could be scanned and made available for public-facing and research purposes.

### 2.1 Data collection

#### Setting

Dreamachine was one of ten major creative projects commissioned for UNBOXED: Creativity in the UK, a UK government-funded festival. It ran in 2022 as a large-scale public installation, presenting a closed-eye stroboscopic light sequence to groups of up to ∼20 or ∼30 people (depending on venue). It was produced by Collective Act Ltd. (CAL), which oversaw venue setup, attendee screening, and the collection and initial filtering of post-session drawings. The installation visited four locations in the UK: Belfast, Cardiff, Edinburgh, and London, delivering approximately 8–10 shows per day to a total audience of over 40,000 attendees. Several of the present authors (DJS, FM, AKS, and TH) provided scientific and philosophical input to the project. The installation was primarily an artistic project, with a secondary role as a platform for collecting observational data outside a controlled laboratory context.

#### Attendee population

More than 40,000 members of the UK public aged 18 or older attended Dreamachine during its 2022 run. Attendees were screened for contraindications to SLS before attending, including self-reported assessment for photosensitivity, epilepsy, migraine, prior adverse responses to flashing lights, and other conditions associated with heightened susceptibility to visually induced seizures or discomfort. Individuals with such sensitivities were directed to a non-SLS version of the event, which accounted for a minority (16.4%) of attendance. Site attendance was distributed across London (42.2%), Edinburgh (22.5%), Belfast (22.5%), and Cardiff (12.8%).

The distribution of scanned drawings is similar, with London contributing 35.7%, Edinburgh 31.4%, Belfast 25.8%, and Cardiff 7.2%, with 0.7% missing location metadata. These differences reflect the fact that not all attendees produced drawings, and that not all drawings produced were subsequently scanned by CAL or retained for release.

#### Stimulus

Dreamachine offered two formats: the High Sensory (HS) format employed SLS, while the Deep Listening (DL) format did not. HS sessions consisted of a 20-minute sequence of closed-eye SLS sequence designed to elicit SIVHs, with varying brightness, frequency, and duty cycle, accompanied by music. SLS was delivered using two ceiling-mounted theatre stroboscopes per attendee. The DL sessions were identical in structure and duration but used non-stroboscopic coloured lights.

#### Drawing Procedure

Following their Dreamachine session, attendees (as a group) entered a shared “reflection space” where they could reflect on their experience in several ways, including by drawing. All activities were entirely optional. Attendees were free to leave at any time.

At a drawing table (Fig. 2), attendees were supplied with black or white A3 paper and coloured dry pastels. Drawings from previous sessions were arranged in the centre of the table (Fig. 2). Attendees were given the option to draw their experience from their Dreamachine session. Attendees could attend multiple Dreamachine sessions and, if they chose, produce multiple drawings.

**Figure 2.**
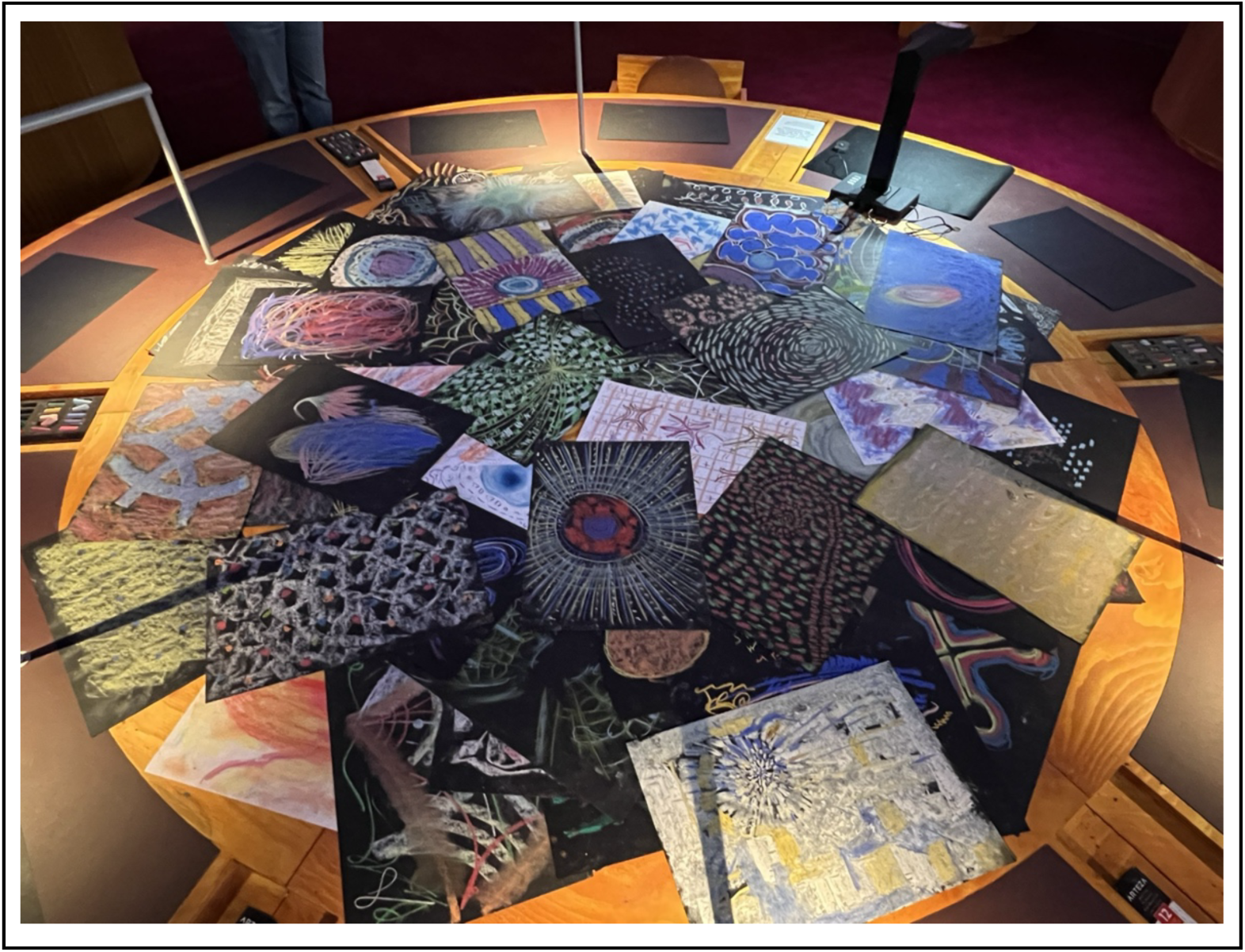
Data collection at Dreamachine. Attendees in the post-session “reflection space” used dry pastels and paper to create drawings representing their experiences following their Dreamachine session.

Attendees who created drawings were also given the chance to scan them, adding them to the dataset. Due to logistical issues with the data pipeline of the Dreamachine, scanned drawings were not labelled with their corresponding session type (HS or DL), limiting interpretability (we return to this in Section 4.2). As the majority of Dreamachine sessions were HS sessions involving SLS (83.6% of attendees), most scanned drawings are expected to predominantly reflect SIVH-related content. Drawings originating from DL sessions, which did not involve SLS, were not expected to contain SIVH-related content and were therefore treated as noise (non-SIVH data) in the analysis.

Because drawings were produced in an open, reflective environment, we cannot definitively determine whether any given drawing directly depicts a simple VH, a complex VH, or non-hallucinatory imagery. These data must therefore be interpreted as indirect, attendee-generated representations of experiences following HS or DL sessions, rather than as ground truth representations of hallucinatory content. Given this mixture, recurring geometric motifs observed across the dataset are interpreted as predominantly reflecting SLS-induced simple VHs, consistent with prior reports of SIVH content (Smythies 1959; Allefeld et al. 2011; Mauro et al. 2015).

### 2.2 Data Analysis

An analysis pipeline was developed to cluster drawings according to their visual features (Fig. 3). Following preprocessing, feature vectors were extracted using a pretrained vision transformer, reduced in dimensionality, and clustered to identify groups of drawings that shared similar visual features. We then conducted a qualitative thematic analysis of the resulting clusters, assigning descriptive codes to each cluster based on its dominant visual characteristics and mapping them onto higher-level categories to support interpretability and to relate the data-driven groupings to existing theoretical frameworks for simple VHs.

**Figure 3.**
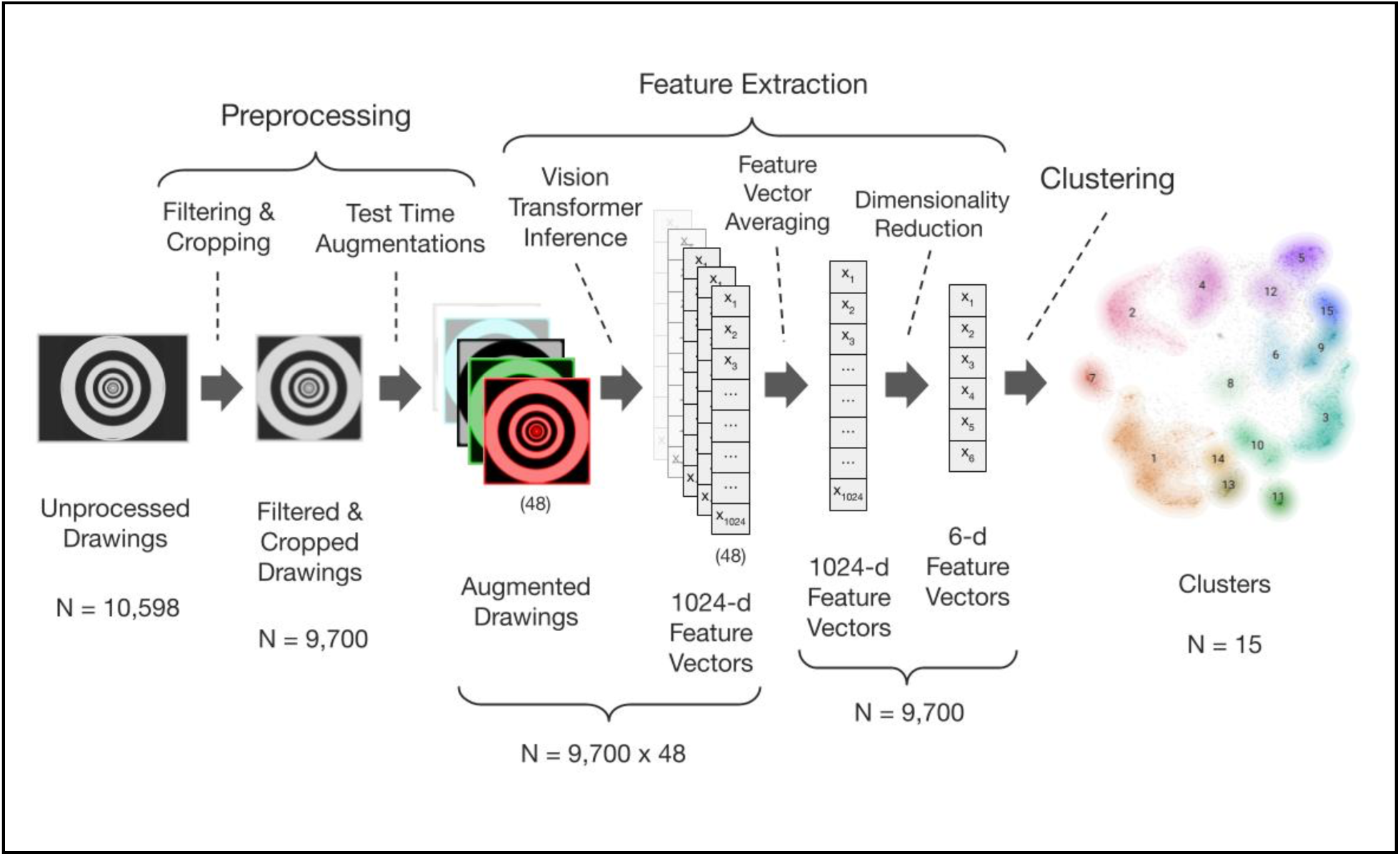
Data processing and clustering pipeline. Starting from 10,598 scanned images, we removed corrupted, duplicate, incorrectly cropped, and text-only images and applied centre-cropping. This yielded 9,700 analysable images. We then applied histogram equalisation and Gaussian smoothing, without further reducing the dataset. Test time augmentations were applied, resulting in 48 augmented drawings for every drawing in the dataset. Each augmented drawing was then embedded using the DINOv2 Vision Transformer (ViT) to obtain a 1,024-dimensional feature vector. The 48 feature vectors of a drawing were averaged, resulting in one feature vector per drawing in the dataset. Dimensionality was first reduced using principal component analysis (PCA; retaining > 90% of the variance) and then using Uniform Manifold Approximation and Projection (UMAP) to a six-dimensional feature space. The resulting six-dimensional embeddings were then clustered using the HDBSCAN algorithm, yielding 15 distinct clusters in the final solution.

#### Preprocessing

The organisers of Dreamachine (CAL) pre-filtered the scans of the drawings (N = 10,598) to remove duplicates, obstructed views, low-quality scans, and explicit content, and to crop the images to the boundaries of the paper, before releasing the dataset to researchers and the public (https://dreamachine.world/drawings/). A subset of images consisted solely of written textual reflections on lined paper; these were excluded from the present analysis. We performed additional preprocessing, excluding corrupted images, remaining duplicate images, and over-cropped or misaligned images, resulting in a final dataset of 9,700 drawings. We centre-cropped images to 95% of the original height, matched width to height, and resized all images to 448 x 448 pixels, the input size required by the DINOv2 model. To reduce capture artefacts such as uneven lighting, shadows, and contrast fluctuations introduced during scanning or photography, we applied local histogram equalisation (exposure/lighting normalisation) and a Gaussian blur (σ = 1, 7×7 kernel) to each image.

To accommodate variations in exposure, lighting, drawing and paper colour, and orientation, we used test-time augmentations (TTA), where multiple transforms of each image are embedded independently and are averaged to produce a more robust and generalisable feature representation. For each included image, we generated 16 transforms (colour inversion, horizontal/vertical flips, rotations 0°, 90°, 180°, 270°), along with random jitter applied to brightness, contrast, saturation, and hue (Fig. S2). To balance coverage and computational cost, we applied three TTA sets per image, producing 48 augmented views per image and a corresponding set of 48 feature vectors. Given the 9,700 images, this resulted in 465,600 augmented images (48 per image), all of which were used as inputs to the ViT (Fig. 3).

#### Feature Vector Extraction

Feature representations were extracted using a Vision Transformer (ViT), a deep learning architecture that captures global image features more effectively than convolutional neural networks (Naseer et al. 2021), making it well-suited for detecting the structured geometric patterns associated with SIVHs. We used DINOv2 ViT-L/14, a self-supervised model developed and trained on ∼142 million naturalistic images by Meta.

The model includes additional learnable ‘register’ tokens (extra vectors appended to the input sequence that the network optimises during training to store latent features), which improve representation quality and feature distillation (Oquab et al. 2024). DINOv2 produces robust, transferable visual representations across a wide range of downstream tasks (Oquab et al. 2024), and its features are designed to capture object parts and scene geometry in a domain-agnostic manner, making it appropriate for analysing abstract geometric drawings. Consistent with this, ViT-based self-attention architectures have been shown to emphasise global shape information and exhibit strong transfer performance (Ericsson et al. 2021; Naseer et al. 2021).

Model inference for the Dreamachine drawing dataset was performed using Google Colab running on an NVIDIA A100 GPU. The inference output included 48 feature vectors per drawing in the Dreamachine dataset, each comprising 1,024 dimensions, which were averaged into a single feature vector of the same length. Averaging embeddings across test-time augmentations has been shown to improve embedding robustness and downstream performance, including classifier accuracy, in related contexts (Ashukha et al. 2021).

#### Clustering

To cluster the feature vectors, we first reduced their dimensionality in two stages and then applied a density-based clustering algorithm. Feature vectors were L2-normalised prior to dimensionality reduction so that Euclidean distances corresponded to cosine similarity–the distance metric used in downstream clustering, which is standard practice for image embedding. We then applied principal component analysis (PCA) to minimise noise unlikely to contribute to meaningful clustering, retaining components explaining ≥90% of the variance (253 components; (Jolliffe 2002)). Next, we applied Uniform Manifold Approximation and Projection (UMAP; (McInnes et al. 2018) to produce a six-dimensional embedding that preserved salient structure while yielding a feature space more suitable for density-based clustering.

In this six-dimensional space, we used the Hierarchical Density-Based Spatial Clustering of Applications with Noise (HDBSCAN; (Campello et al. 2013)), which identifies clusters from a hierarchy of density-based partitions by selecting the most stable structures and labels atypical points as noise rather than assigning all drawings to clusters. Unlike K-means, HDBSCAN does not assume spherical clusters, and unlike DBSCAN it can accommodate clusters of varying density, properties well-suited to heterogeneous, unconstrained drawings.

Parameters for UMAP and HDBSCAN were selected using a combination of *a priori* choices, systematic grid search, and qualitative assessment of cluster structure and interpretability (see Supplementary Materials). The Density Based Clustering Validity (DBCV) metric, a measure of clustering quality, was used to exclude candidate solutions with low values (Fig. S3). Candidate solutions were compared qualitatively to reduce the proportion of data points classified as noise (unclustered), obtaining an interpretable yet sufficiently granular number of clusters, and ensuring clear partitioning of visual features between clusters as visualised in the two-dimensional UMAP projection.

#### Qualitative analyses

We conducted a two-stage inductive thematic analysis, in which patterns are derived directly from the data rather than imposed by predefined categories. First, two of the authors (EG and TH) inspected all the drawings within each of the 15 clusters and iteratively generated descriptive codes that captured the dominant visual regularities in each cluster (e.g., “spirals”, “radial lines”; see Supplementary Materials for the complete image set by cluster; Fig. 4).

**Figure 4.**
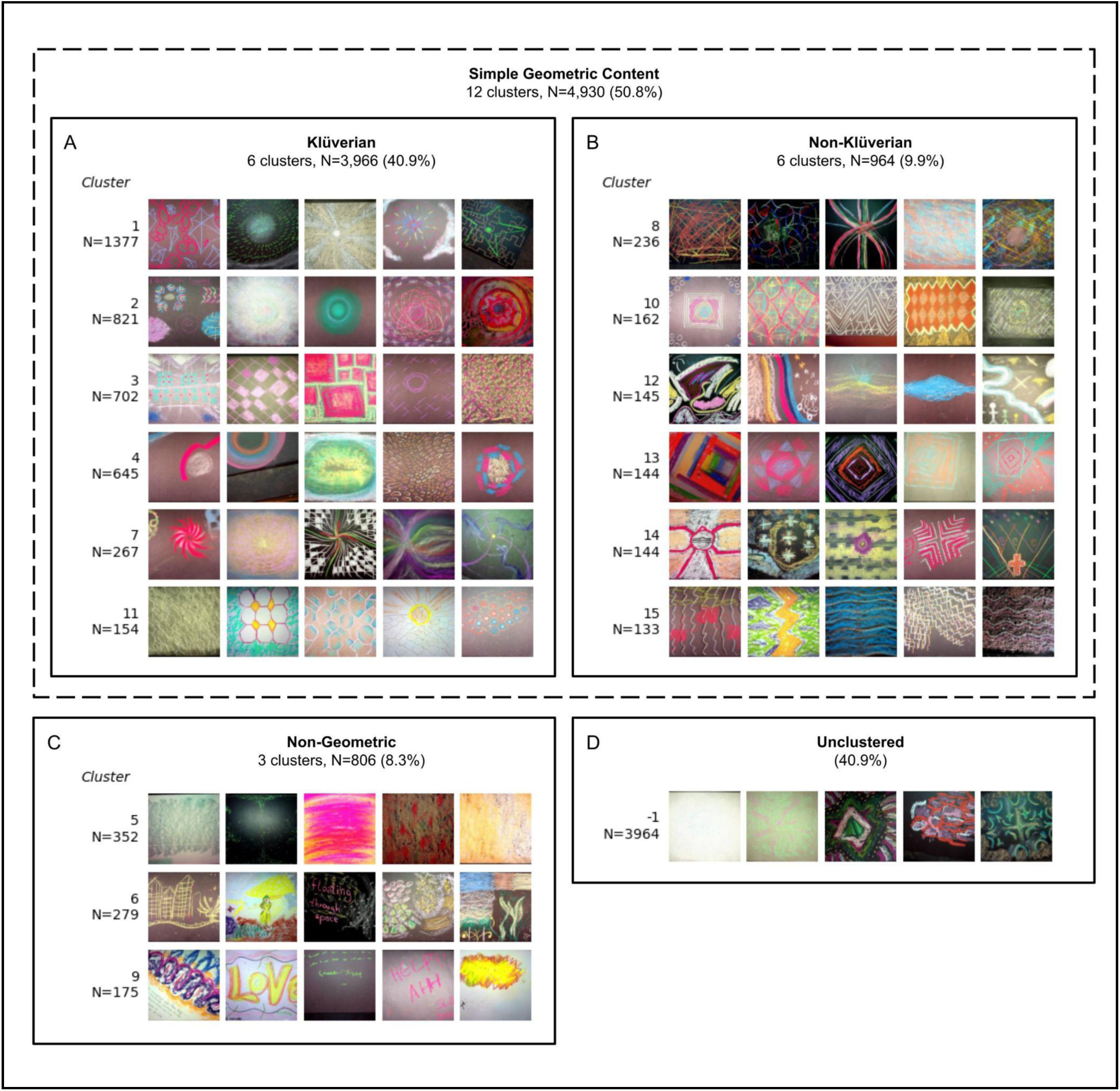
Overview of clustering solution and thematic categories. Clusters identified by HDBSCAN are grouped into three higher-level categories derived from qualitative thematic analysis: (A) *Klüverian* simple geometric content, (B) *Non-Klüverian* simple geometric content, and (C) *Non-Geometric* content (green), plus (D) the Unclustered drawings. The *Klüverian* and *Non-Klüverian* panels together form the “Simple Geometric Content” category. Within each panel, each row corresponds to a single cluster and shows representative drawings randomly sampled from that cluster. Labels to the left of each row indicate the cluster ID (top) and number of drawings in the cluster (bottom); cluster IDs are ordered by decreasing cluster size (Cluster 1 is largest). Drawings in the Unclustered panel (cluster ID – 1) were not assigned to any cluster by HDBSCAN. The cluster codes are: (–1) Unclustered (HDBSCAN noise); (1) Radial lines, repeating rotationally symmetric patterns; (2) Concentric circles, spirals; (3) Lattices, checkerboards; (4) Circles; (5) Uniform colours, diffuse coloured forms; (6) Miscellaneous geometries, semantic content; (7) Spirals; (8) Intersecting curved lines; (9) Scribbled lines, writing; (10) Diamonds, triangles; (11) Hexagonal lattices; (12) Curved lines; (13) Concentric diamonds/squares; (14) Crosses; (15) Wavy/zig-zag stripes.

Second, these codes were compared across clusters and consolidated into three higher-level categories, guided by the Klüver form-constant framework while allowing for data-driven extensions (see Supplementary Materials). Coder disagreements were resolved by discussion to reach consensus coding for each cluster.

## 3. Results

Based on qualitative and quantitative evaluations of the candidate clustering outcomes—including assessments of cluster quality (DBCV), interpretability, visual separability, proportion of noise, and overall cluster count—a final clustering solution was reached in which 59.1% (N = 5,736) of the images were assigned to 15 clusters, with the remainder left unassigned by HDBSCAN. Qualitative thematic analysis revealed that 12 clusters predominantly contained geometric formations, of which six corresponded to the well-documented Klüver form constants (henceforth “Klüverian”), and six comprised additional, systematically recurring geometric formations (“non-Klüverian”). The remaining three clusters captured non-geometric content such as diffuse colour fields and semantic or symbolic visual content. Figure 4 shows an overview of clusters and their thematic categories.

### 3.1 Selection of the Clustering Solution

The selected clustering solution classified 59.1% of drawings into 15 clusters, with the remaining 3,964 (40.9%) drawings left unclustered (designated noise by HDBSCAN). This proportion of unclustered images is consistent with expectations for a heterogeneous, unconstrained dataset containing drawings with highly idiosyncratic content, mixtures of geometric and non-geometric elements, and contributions from both HS and DL sessions. This clustering solution was selected from a grid search over HDBSCAN’s *min_samples* parameter (2-100), which controls the conservativeness of the clustering solution (i.e., the number of clusters and the proportion of noise). Candidate solutions spanned 8-19 clusters with 45.8-26.8% noise (Fig. S3). Visual inspection of the two-dimensional UMAP projection (Fig. 5) indicated that clusters captured salient and distinct visual features, balancing interpretability and granularity.

**Figure 5.**
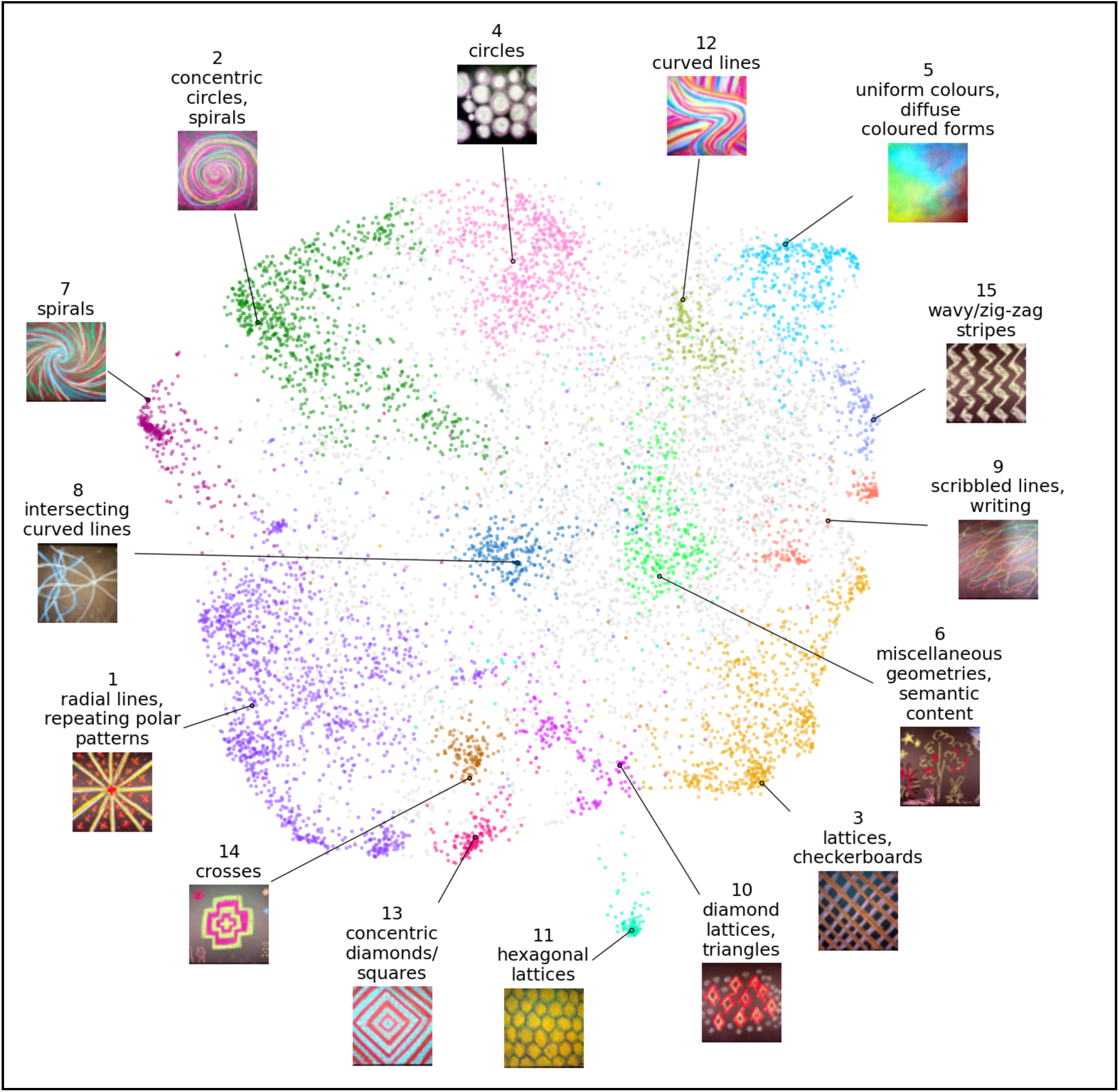
Two-dimensional UMAP projection of the Dreamachine image embeddings clustered using HDBSCAN. Each point represents a single drawing, projected into two dimensions using UMAP for visualisation purposes. Points are coloured according to HDBSCAN cluster membership, with grey points indicating drawings not assigned to any cluster. For each of the 15 clusters, a representative drawing (selected for visual clarity) and its descriptor are displayed to illustrate the characteristic visual features captured by that cluster. Cluster numbers correspond to those reported in Fig. 4. An interactive version of this visualisation is available online at: https://ejgrove.github.io/dreamachine_visualization/

Given the absence of a single objective criterion for an “optimal” solution, the grid search produced a family of plausible clustering solutions, each trading off cluster number against noise proportion. We selected the 15-cluster solution because it combined moderate noise levels with visually coherent, well-separated clusters, providing a representative and interpretable overview of the dataset’s geometric structure.

### 3.2 Clustering Solution

Cluster sizes for the 15 HDBSCAN clusters (excluding the unclustered group) ranged from 133 to 1,337 drawings (mean = 382; SD = 356). The Density Based Clustering Validity (DBCV) metric – ranging from 0 to 1 – was 0.38, indicating moderate density–separation structure for this clustering solution. Alternative solutions had higher DBCV values (>0.5); however, the final clustering solution reported here was selected by jointly optimising internal cluster-validity metrics and qualitative interpretability, balancing cluster structure, the proportion of drawings left unclustered, and the overall number of clusters (Fig. S3).

There was a statistically significant association between cluster and Dreamachine location χ² = 192.4, df = 60, p < 0.001; Supplementary Fig. S4). However, the effect size was negligible (Cramér’s V = 0.072), indicating only minor deviations from uniformity and that clusters were broadly similar in their geographic distributions.

### 3.3 Thematic analysis of clusters

Clusters were visually inspected to characterise their shared visual features and assigned to one of three thematic categories, *Klüverian*, *Non-Klüverian*, and *Non-Geometric*, shown in Fig. 4. Thematic categories were derived through inductive thematic analysis, guided, but not constrained, by the Klüver form-constant framework. The *Klüverian* and *Non-Klüverian* categories comprise of geometric formations, with the former containing the four Klüver form constants and the latter comprising systematically recurring geometric patterns that are not captured by the Klüver framework. The *Non-Geometric* category includes non-geometric visual content, such as semantic imagery, text, and diffuse or unstructured colour fields. Unclustered drawings were not assigned to any thematic category.

#### Simple Geometric Hallucinations

Two categories, *Klüverian* and *Non-Klüverian*, comprise the simple geometric content that is observed across much of the Dreamachine dataset, containing 50.8% of all drawings and 85.9% of the clustered subset.

#### Klüverian

The *Klüverian* category consisted of six clusters (N=3,966, 40.9%) containing geometry that corresponded to one or more of Klüver’s form constants. Three clusters (1, 2, 7) were dominated by rotationally symmetric geometries, including tunnel, spiral, and cobweb form constants: Cluster 1 “radial lines, repeating rotationally symmetric patterns” [N=1,377, 14.2%; tunnel, cobweb], Cluster 2 “concentric circles, spirals” [N=821, 8.5%; tunnel, spiral], and Cluster 7 “spiral” geometries” [N=267, 2.8%; spiral]). One cluster (Cluster 3 “lattices, checkerboards” N=702, 7.2%), was characterised by translationally symmetric lattice patterns. Two clusters contained mixed translational and rotational symmetries (Cluster 11 “hexagonal lattices” [N=154, 1.6%; lattice, tunnel] and Cluster 4 “circles” [N=409, 4.2%; lattice, tunnel]). Across the Klüverian category, all four of Klüver’s canonical form constants were identified: tunnel (Clusters 1, 2, 4, 11), spiral (Clusters 2, 7), cobweb (Cluster 1), lattice (Clusters 3, 4, 11) (see Fig. 6A for schematic illustrations).

**Figure 6.**
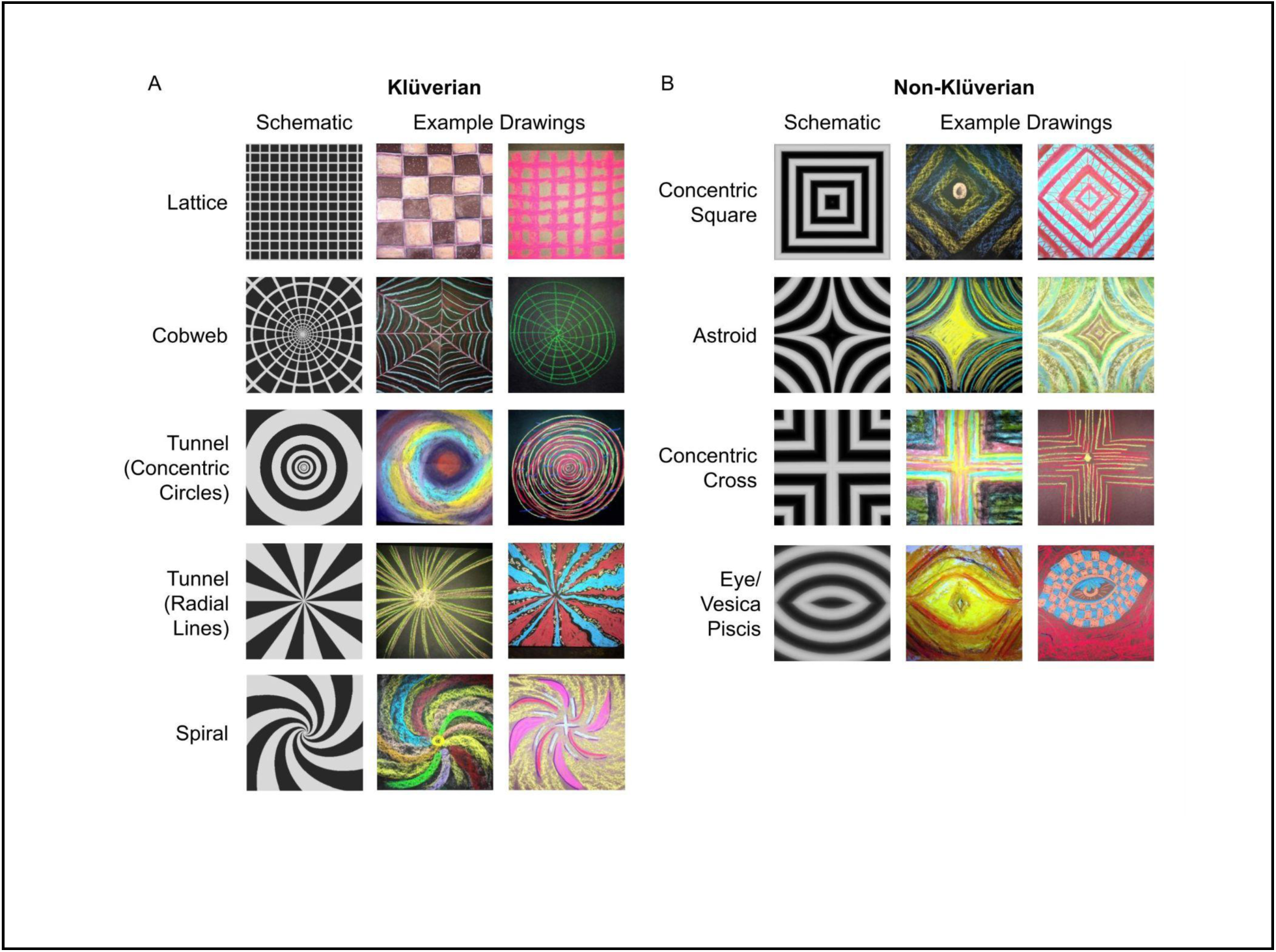
Geometric motifs of SIVHs: Klüver form constants and non-Klüverian extensions. For each subtype, a schematic is shown on the left and representative attendee drawings on the right. (A) Klüver form constants; (B) Non-Klüverian geometries, including the vesica piscis (the intersection of two equal-radius circles whose centres lie on each others’ circumference). Schematics were generated using a custom computational hallucination-recreation interface developed by author TH. Attendee drawings are reproduced with permission from Collective Act Ltd.

#### Non-Klüverian

Six clusters (N = 964; 9.9%) were classified as *Non-Klüverian,* comprising systematically recurring geometric patterns not captured by Klüver’s form constants. An overview of these forms is shown in Fig. 6. Two clusters exhibited pronounced fourfold rotational symmetry: Cluster 13 (“concentric diamonds/squares”; N = 162; 1.5%) and Cluster 14 (“crosses”; N = 144; 1.5%). Cluster 10 “diamond lattices, triangles” (N=162, 1.7%) contained lattice-like structure, but was distinguished by concentric triangular motifs, which formed a spatially distinct cluster in the UMAP projection (Fig. 5), and, to our knowledge, have not previously been reported in the SIVH literature. Cluster 15 (“wavy/zig-zag stripes”; N=133, 1.4%) comprised undulating, stripe-like geometries. Finally, Cluster 8 (“intersecting curved lines”; N = 236, 2.4%) and Cluster 12 (“curved lines”; N = 145, 1.5%) contained diverse but related patterns characterised by overlapping arcs, flowing contours, and curvilinear structure.

#### Non-Geometric

The *Non-Geometric* category consisted of three clusters accounting for 8.3% of the drawings. Cluster 5 was labelled “uniform colours, diffuse coloured forms” (N = 352, 3.6%), and comprised drawings characterised by blended or overlapping colour regions without clear geometric structure or semantic imagery. Cluster 9 “scribbled lines, writing” (N = 175, 1.8%) contained written text and loosely scribbled marks, which were visually similar to one another but did not form clear geometric patterns. Cluster 6 “semantic content” (N = 279, 2.9%), included drawings depicting recognisable objects, symbols, or scenes, and may reflect complex visual hallucinations or post-session reflective imagery rather than simple geometric hallucinations.

#### Unclustered

Around 40.9% (N=3,964) of drawings were not assigned to any cluster by HDBSCAN. These drawings contained idiosyncratic visual motifs that did not recur with sufficient numbers to form a single cluster or combined features characteristic of multiple clusters. Despite this heterogeneity, some recurring motifs–present at frequencies below the clustering threshold–were observed within the unclustered set, including some less common geometric patterns and realistic drawings of eyes (Fig. 6B).

## 4. Discussion

Using a data-driven computer-vision approach applied to a large set of attendee drawings produced following stroboscopic light stimulation (SLS), we show that the phenomenology of simple visual hallucinations (VHs) encompasses a broader and more structured range of geometric forms than is captured by classical form-constant taxonomies. While many drawings align with Klüver’s form constants, a substantial proportion exhibit systematically recurring non-Klüverian geometries, including translationally symmetric and fourfold-rotationally symmetric patterns. These observations place empirical constraints on current neural–field models of simple VHs, which primarily account for rotationally symmetric percepts, and highlight the value of large-scale, participatory datasets analysed with modern computer-vision methods as a complementary, hypothesis-generating route to quantitative phenomenology.

### 4.1 Visual Phenomenology of SIVHs: Theory and Observation

The majority of the data (12 out of 15 clusters) were dominated by drawings of geometric patterns, indicating that simple geometric content predominated in this setting. This is consistent with controlled laboratory work showing that SLS reliably elicits simple geometric VHs (Mauro et al. 2015; Allefeld et al. 2011; Hewitt et al. 2025). The non-geometric images in the dataset likely reflect multiple contributing factors. For example, diffuse colour experiences (which constitute the majority of drawings in Cluster 5) align with previous reports of chaotic or unstructured SIVHs (Beauté et al. 2025; Smythies 1959), but may also reflect contributions from Deep Listening (DL) sessions, which lacked SLS and instead emphasised music and ambient coloured light.

The semantic drawings and textual content in Clusters 9 and 6, by contrast, likely arise from a mixture of hallucinatory and non-hallucinatory sources. Some of these drawings are consistent with complex VHs, which are occasionally reported in SIVHs (Schwartzman et al. 2019; Bartossek et al. 2021; Beauté et al. 2025; Geiger 2004), whereas others likely represent conceptual or reflective responses that do not directly depict an attendee’s perceptual experiences in the Dreamachine. Finally, the unclustered drawings contained heterogeneous mixtures of geometric and semantic elements that did not cohere into a single motif, indicating that in large-scale, naturalistic settings such as Dreamachine, a substantial proportion of participant-generated drawings cannot be cleanly partitioned into discrete geometric classes, even when produced under broadly similar stimulation conditions.

Among the observed geometric formations, a large portion of the images consisted of Klüver’s form constants (Fig. 6A), while a substantial subset were non-Klüverian in nature, such as concentric squares, crosses, and astroid patterns (rounded, pinched-square curves) (Fig. 6B). A notable example that places constraints on key assumptions of dominant neural-field models of simple VHs is the prevalence of fourfold rotational symmetry, defined by repeated rotational structure every 90°. All images shown in Fig. 6B exhibit this property. Clear fourfold rotational symmetry appeared across multiple clusters, including those dominated by concentric diamonds or squares (Cluster 13) and crosses (Cluster 14). Concentric squares, crosses, and lattices have been noted in some earlier SLS case studies (Smythies 1959; Allefeld et al. 2011). However, their prevalence has not been quantified. The frequency of this symmetry (approximately 5.8% of the *Simple Geometric Content* drawings and 3% of all drawings) suggests that fourfold rotationally symmetric patterns constitute an under-characterised subclass of simple VHs. Within this subclass, astroid shapes constitute an additional motif that, to the best of our knowledge, has not previously been reported in accounts of SIVHs. Importantly, the novel and under-reported forms shown in Fig. 6 were each observed repeatedly across all four Dreamachine sites, arguing against site-specific or procedural artefacts.

From a mechanistic perspective, the fact that regular, structured simple VHs recur under relatively simple bottom-up stimulation in a large, naturalistic sample, and are also reported across aetiologically distinct hallucinatory conditions, suggests an important contribution from the architecture and dynamics of early visual cortex.

The observed geometries place new constraints on neural-field models of simple VHs. Current retinocortical neural-field models that apply a log-polar mapping from cortical coordinates to the visual field can reproduce many rotationally symmetric geometries, including spirals, tunnels, and cobweb-like Klüver form constants. However, these models do not readily account for the broader range of geometries observed here: after the log–polar transform, most cortical pattern families map onto rotationally symmetric percepts, leaving translational and fourfold-symmetric formations poorly accounted for. In particular, lattice-like structures, concentric squares, crosses, and astroids are not readily generated within current formulations of these models.

Extending models to reproduce the empirically observed range of VH geometries has the potential to advance our understanding of the mechanisms by which early visual cortical architecture and dynamics shape conscious experience, veridical and hallucinatory alike, by more explicitly linking the organisation and dynamics of early visual cortex to the structured patterns observed in SIVHs.

### 4.2 Limitations

Interpretation of our findings is constrained by both the nature of the dataset and the exploratory design of the analysis. Drawings were collected during a public art-science installation without experimental control, explicit instructions on how (or indeed whether) to depict SLS experiences, or verification that attendees underwent the High Sensory (HS, SLS) as opposed to the Deep Listening (DL, non-SLS) experience. The absence of HS/DL labels for individual drawings is a key limitation of the dataset. We investigated several approaches to retrospectively apply labels (e.g., by matching scan timestamps to recorded session details); however, these efforts were unsuccessful. Several factors mitigate this problem. First, many more participants underwent the HS (83.6%) compared to the DL (16.4%) experience. Second, it is possible that a lower proportion of DL attendees (compared to HS attendees) made drawings, given the reduced emphasis on visual experience in the DL condition, in which case the proportion of DL drawings would be even less than 16.4%. Altogether, the DL drawings are best thought of as additional ‘noise’ within the dataset – potentially populating the unclustered part of the dataset. Nonetheless, the presence of DL drawings in the dataset constrains inference about SLS-specific effects and should be borne in mind when interpreting the results.

Although we refer to “simple VHs” for continuity with the literature, and because SLS induces such experiences, it is important to note that the present analysis concerns drawings produced after Dreamachine sessions rather than direct, immediate, trial-by-trial reports of hallucinatory experiences. The Dreamachine environment introduced several possible confounds, including the open and socially interactive setting, the influence of attendees viewing other attendees’ drawings, conversations between attendees, and possible demand characteristics, whereby attendees might (implicitly or explicitly) attempt to reproduce what they believed they were “supposed” to see. Any or all of these factors may have shaped the resulting drawings.

Moreover, the drawings represent retrospective reconstructions of remembered percepts and are further constrained by individual differences in drawing ability, as well as variation in how comfortable or inclined individuals were able to depict abstract geometric versus semantic content.

Our unsupervised computer-vision pipeline provided a data-driven method to group drawings by their shared visual features. However, the clustering solution was nevertheless influenced by methodological choices, including the selection of the vision model, feature extraction steps, clustering algorithm, and the subjective evaluation of cluster interpretability. Different parameter settings could yield alternative clustering solutions. A related approach was taken by Beaute et al., who clustered free-text reports from Dreamachine visitors using UMAP and HDBSCAN, but were able to optimise their solution using language-based coherence metrics and to label clusters using a large language model. In contrast, no comparable quantitative metrics currently exist for evaluating the interpretability of visual geometric clusters, necessitating qualitative judgement in the present study. Consistent with this constraint, we observe finer-grained substructures are visible in Cluster 1 (e.g., astroids and “radial lines, repeating rotationally symmetric patterns” within Cluster 1). While the present analysis sought to broadly characterise the range of geometric forms commonly observed in simple VHs, further hierarchical clustering solutions could produce a more detailed description of classes and sub-classes of hallucinatory geometric formations.

Altogether, our analysis should be viewed as an interpretable and practically useful partition of the dataset rather than a unique or definitive solution. In addition, the absence of trial-by-trial linkage between drawings and specific SLS parameter combinations (e.g., frequency or brightness) prevents direct inference about parameter–phenomenology relationships. We therefore frame our findings as exploratory and hypothesis-generating, rather than hypothesis-confirming. Despite these limitations, the recurrence of visually coherent clusters across a large, heterogeneous sample suggests the presence of shared perceptual tendencies across attendees, providing a robust empirical foundation for future targeted experimental work.

## 5. Conclusion

We developed a novel data-driven pipeline to examine drawings made following hallucination-inducing exposure to stroboscopic light stimulation. Our results indicate that the range of geometric formations in simple VHs is broader than that implied by the existing accounts, encompassing both classical Klüver form constants and additional, systematically recurring but under-reported geometric motifs. These findings place new constraints on neural-field models of simple VHs, which currently account for a limited subset of the observed patterns. Altogether, our study demonstrates the promise of large-scale participatory approaches to quantitative phenomenology, combining participant-generated drawings with unsupervised computer-vision analysis. Such approaches offer significant potential to constrain and refine mechanistic theories of visual hallucinations, and in turn inform broader accounts of vision, consciousness, and their altered states.

## Acknowledgements

D.J.S. is supported by a Medical Research Council Grant UKRI083. A.K.S. is supported by European Research Council Advanced Investigator Grant 101019254. T.H. is supported by the Margaret Boden PhD scholarship from the University of Sussex. F.M. was supported by a grant for Project SENSOR from the AHRC/UKRI, grant number: AH/Y007638/1. F.M., T.H., D.J.S., and A.K.S. were partly supported by a grant from the UK Govt for their participation in the Dreamachine Programme, as part of Unboxed2022. All authors are grateful to Collective Act Ltd (CAL) and especially to CAL Director Jennifer Crook for making Dreamachine happen and collecting the data on which this paper is based. Note that T.H., F.M., A.K.S., and D.J.S. are collaborators on Collective Act’s Dreamachine programme and received funding from the UK Government/Unboxed 2022 via Collective Act Ltd to support this collaboration.

## 6.0 Supplementary materials

### 6.1 Retinocortical Transforms

Neural-field models of simple visual hallucinations simulate patterns of activity in the early visual cortex. The visual field is mapped onto the cortex through an approximately log–polar retinocortical transform (Fig. S1); therefore, these models invert this mapping to generate predicted visual experiences from simulated cortical activity.

**Figure S1:**
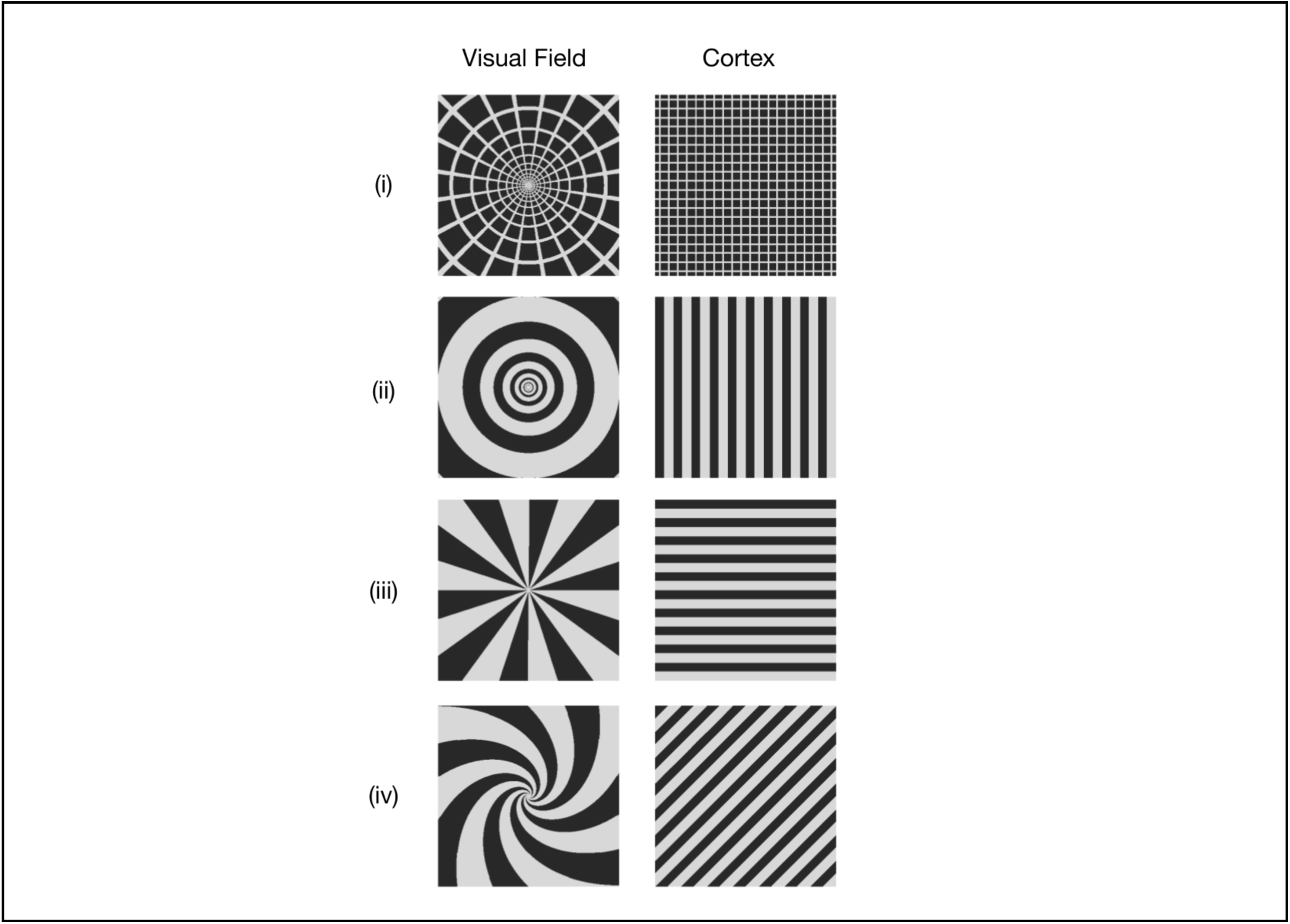
Illustrative retinocortical transforms linking cortical patterns to Klüver form constants. Each row shows a model cortical pattern (right column, Cortex) and the corresponding predicted visual-field percept after applying the inverse retinocortical (log–polar) mapping (left column, Visual Field). Rows (i–iv) illustrate, respectively, the cobweb, tunnel (two variants), and spiral Klüver form constants.

### 6.2 Additional information: dataset

#### Image Filtering and Cropping

Along with the drawings, the same scanning system also captured written textual reflections on Dreamachine, which were identified and removed from the dataset using a convolutional neural network (CNN) trained to distinguish handwritten text from drawn content. Duplicate images were flagged and removed by analysing pairwise cosine similarity scores (> 0.7) between DINOv2 feature vectors. Additional duplicates were manually identified and removed during subsequent work with the dataset. Each duplicate image was visually inspected, and one version was retained based on image quality, orientation, and centrality of the paper within the scan.

Cropped images were validated against the expected dimensions of A3 paper (aspect ratio 1:√2, width approximately 4165 px, height approximately 2945 px). Images that were portrait-oriented or had dimensions less than two-thirds of the calculated paper size (<⅔ × 2945 px height or <⅔ × 4165 px width) were excluded, because these scans typically represented incorrectly captured or heavily cropped pages.

Height was centre-cropped to 2797 px (5% below the average height) to account for variation in paper placement and borders. The width was then cropped to match the adjusted height, producing square images suitable for downstream processing and feature extraction.

**Figure S2:**
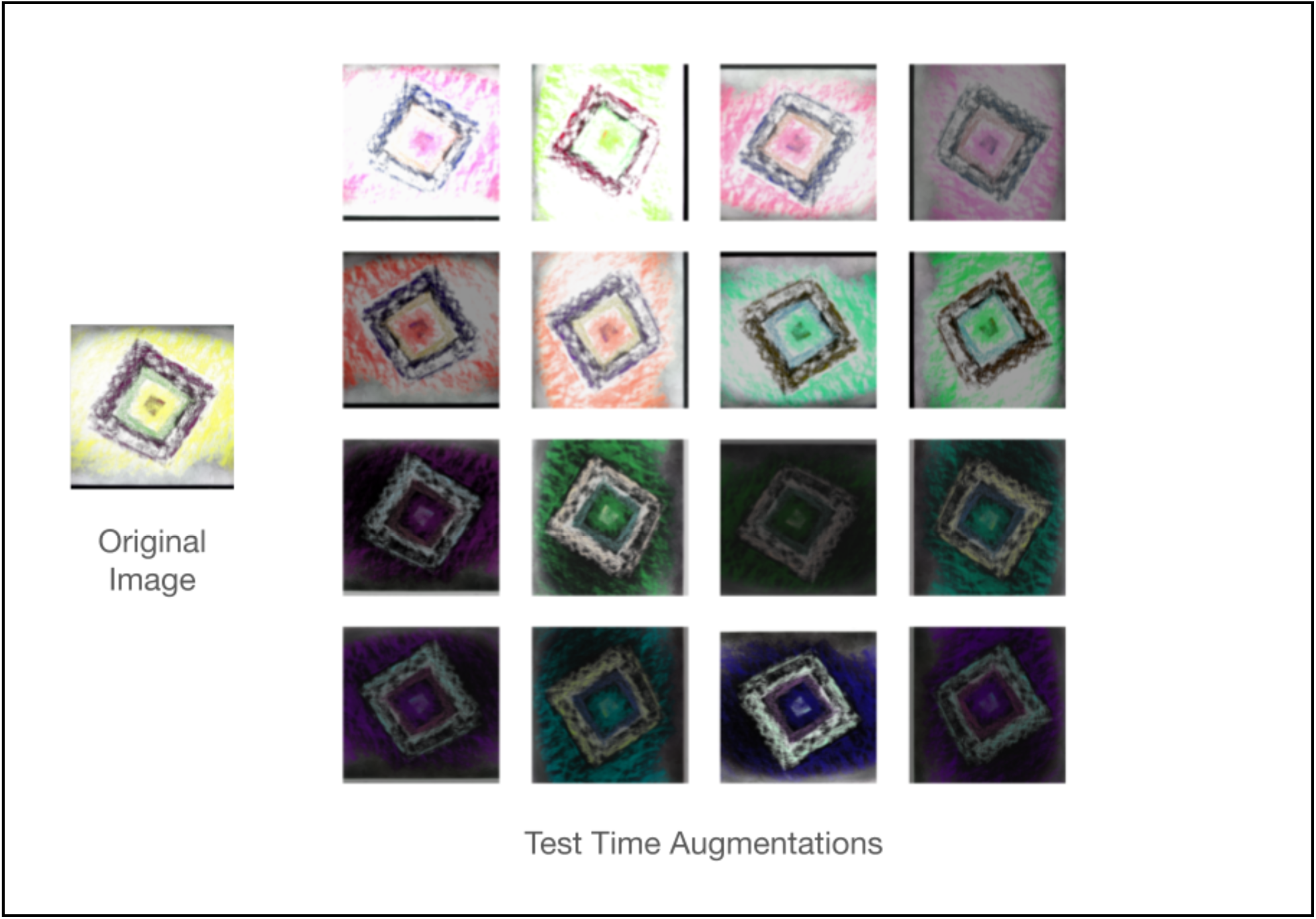
Test-time augmentations applied to a single Dreamachine drawing. The panel on the left shows an example original drawing. The grid on the right illustrates a subset of the 16 test-time augmentations (TTAs) generated for each image, including colour inversion, horizontal and vertical flips, rotations (0°, 90°, 180°, 270°), and random jitter in brightness, contrast, saturation and hue. For each drawing, three sets of TTAs were applied (48 augmented views in total), and the corresponding DINOv2 feature vectors were averaged to obtain a single, more robust embedding for clustering.

### 6.3 Additional information: clustering analysis

#### UMAP

Cosine similarity was selected as the distance metric, consistent with common practice for image embeddings and with DINOv2. The minimum distance parameter was set to 0.0, allowing for compact clusters. We performed a grid search to determine the optimal embedding dimensionality based on the trustworthiness metric, which measures how well local neighbourhood structure from the original feature space is preserved. Trustworthiness plateaued above six dimensions; therefore, we selected a six-dimensional embedding. The *n_neighbors* parameter was selected based on the expected scale of clusters defined by the HDBSCAN *min_cluster_size* (100). Because *min_cluster_size* was set to 100, n_neighbors should be of a similar order of magnitude to ensure that UMAP preserves neighbourhood structure at this scale. This provided a principled rationale that restricted the parameter search space, appropriate for an exploratory rather than purely optimisation-focused analysis.

#### HDBSCAN

The *min_cluster_size* parameter was fixed at 100, corresponding to the smallest cluster size we judged to be interpretable. The *min_samples* parameter was varied from 2-100. The default behaviour sets *min_samples* parameter equal to *min_cluster_size*, however, smaller values are commonly used in practice, so we explored the full range from 2-100 (Fig. S3).

### 6.4 Additional information: qualitative analysis

#### Coding

Descriptions of each cluster’s shared features were developed by manually reviewing the complete set of drawings to identify recurring visual characteristics. Each description emphasises features consistently observed either across an entire cluster or within clearly identifiable subsets. Subsets were defined when distinct shared features were recognised within a cluster during manual inspection. The descriptions of shared features from distinct subsets are separated by commas.

**Figure S3.**
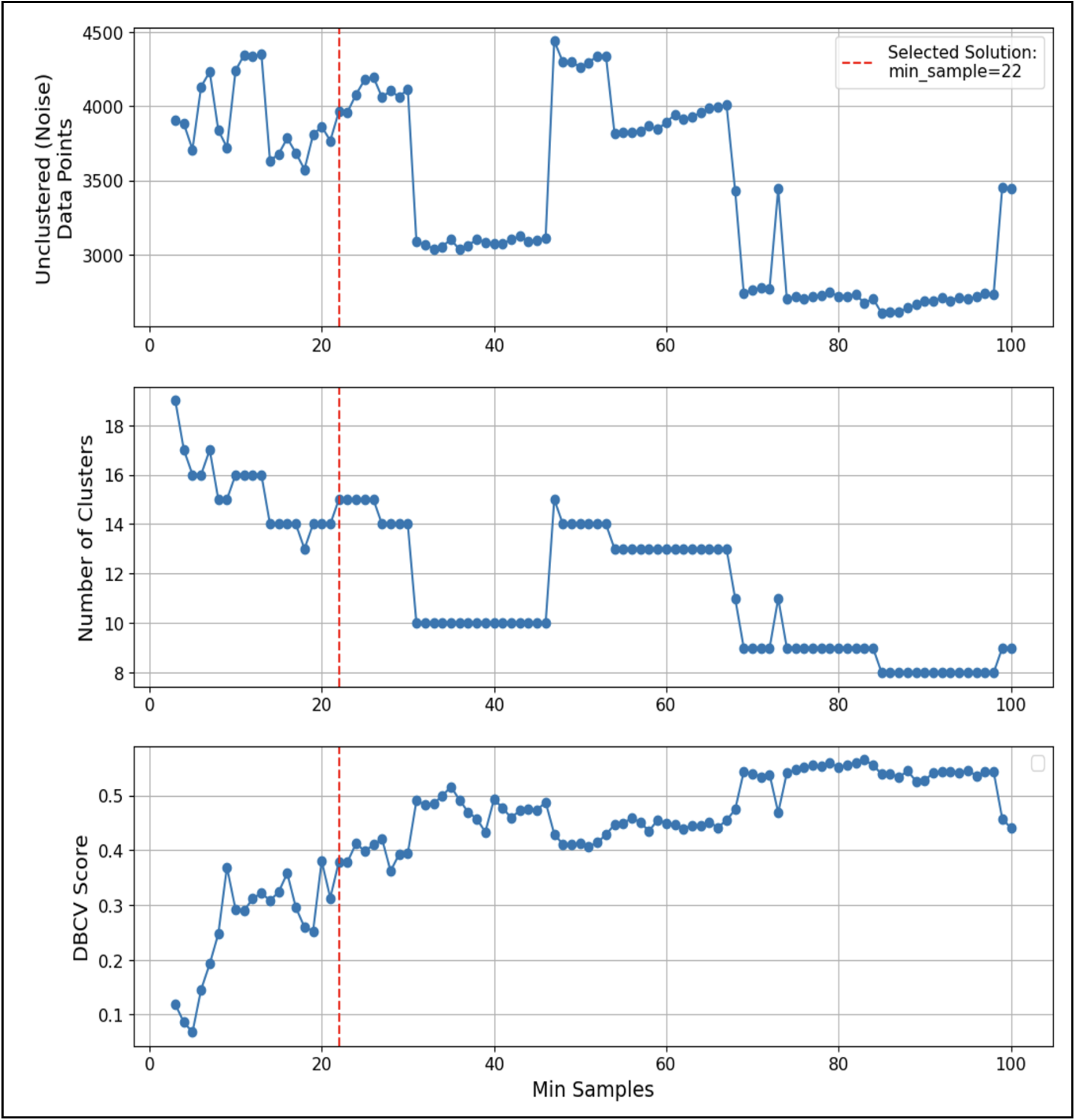
Effects of the HDBSCAN min_samples parameter on clustering solutions. For each candidate value of min_samples (x-axis), the panels show (top) the number of drawings classified as noise (unclustered), (middle) the number of clusters, and (bottom) the Density Based Clustering Validity (DBCV) score, which summarises clustering quality (0–1; higher values indicate more compact, well-separated clusters). The selected solution used min_samples = 22 (red dashed line), balancing noise proportion, cluster count, and DBCV score.

**Figure S4:**
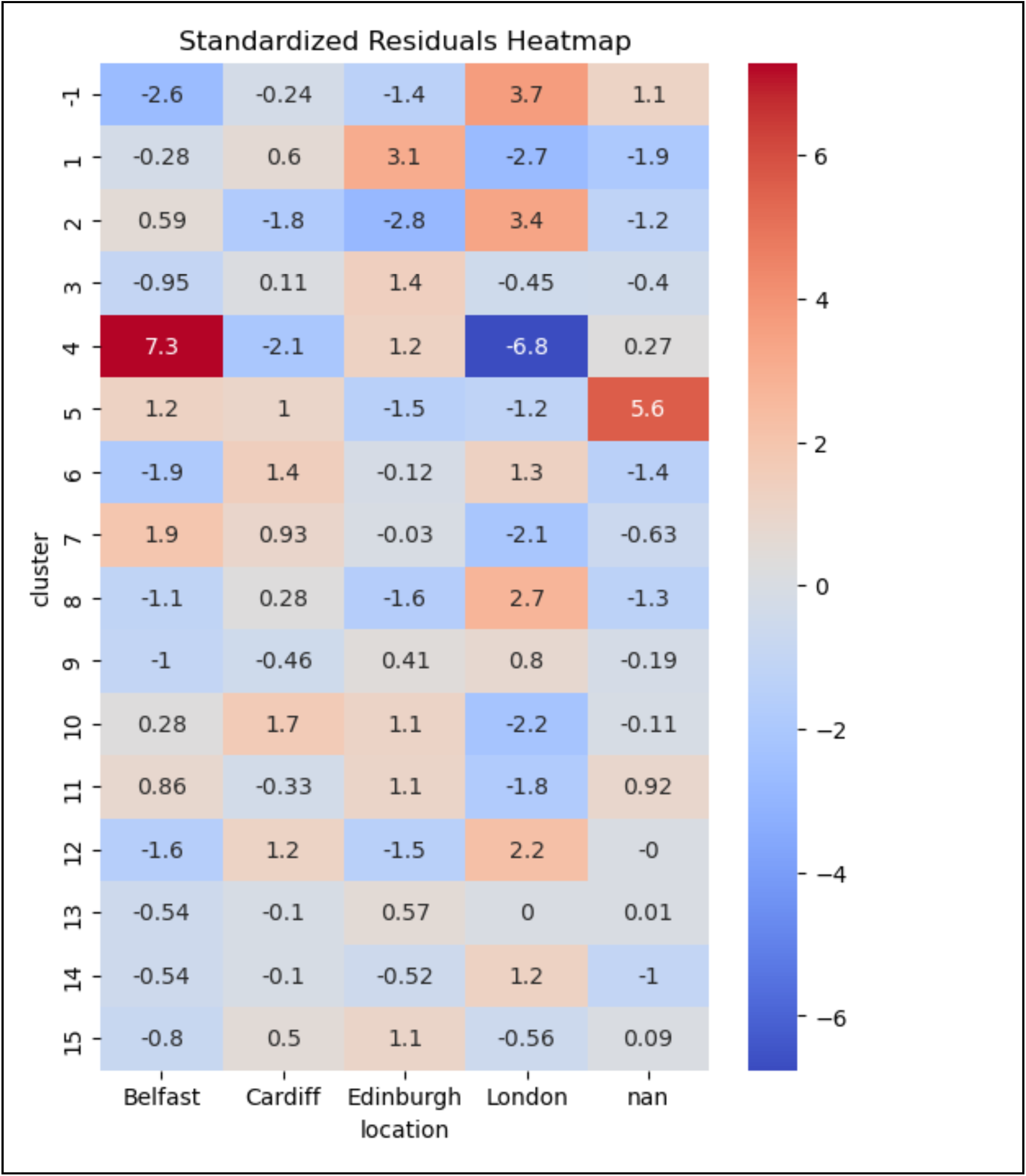
Cluster–location associations across Dreamachine sites. Heatmap of standardised Pearson residuals from a chi-squared test of independence between HDBSCAN clusters and Dreamachine locations. Positive residuals indicate cluster–location combinations that occur more often than expected by chance, and negative residuals indicate combinations that occur less often. Cluster 4 from London occurs significantly less often than expected, whereas Cluster 2 from London, the unclustered images from London, Cluster 4 from Belfast, and Cluster 5 with no location data occur significantly more often than expected. A Bonferroni-corrected critical value of ±3.38 was used to determine statistical significance.

#### Cluster Categories

After coding the cluster descriptors, we applied thematic mapping: all 15 clusters’ descriptors were compared and placed into three categories anchored to Klüver’s form constants: Klüverian geometries, Non-Klüverian geometries, and Non-Geometric drawings.

The interpretation of the Klüver form constants we use in this analysis is defined as (i) *lattice*: translationally symmetric geometries; (ii) *cobweb*: patterns with hybrid radial and concentric rotationally symmetric geometries; (iii) *tunnel*: concentric, radial, and repeating elements with rotationally symmetric geometries; (iv) *spiral*: one or more radial spokes revolving around a central point and extending outward, with curves or angular steps. Klüver’s original description, however, is sufficiently broad that many additional versions of these patterns could qualify.

Here, we distinguished Klüverian from non-Klüverian geometries to reflect both historical usage and current modelling practice, and included only the classical form constants within the Klüverian category.

Clusters dominated by the canonical spirals, tunnels, lattices, or cobwebs were grouped as Klüverian geometries. Clusters with geometries that fell outside the Klüver classification were labelled non-Klüverian geometries (e.g., concentric crosses, diamonds). Some clusters contained both Klüverian and non-Klüverian geometries; these were classified as non-Klüverian. Clusters with specific descriptors, which included semantic, written, or non-geometric content, were classed as non-geometric drawings.

